# Social behavior in bees influences the abundance of *Sodalis* (Enterobacteriaceae) symbionts

**DOI:** 10.1101/192617

**Authors:** Benjamin E. Rubin, Jon G. Sanders, Kyle M. Turner, Naomi E. Pierce, Sarah D. Kocher

**Author notes:** Co-senior authors (Naomi E. Pierce, Sarah D. Kocher).

## Abstract

Social interactions can facilitate transmission of microbes between individuals, reducing variation in gut communities within social groups. Thus, the evolution of social behaviors and symbiont community composition have the potential to be tightly linked. We explored this connection by characterizing the diversity of bacteria associated with both social and solitary bee species within the behaviorally variable family Halictidae using 16S amplicon sequencing. Contrary to expectations, we found few differences in bacterial abundance or variation between social forms, and most halictid species appear to share similar gut bacterial communities. However, several strains of *Sodalis,* a genus described as a symbiont in a variety of insects but yet to be characterized in bees, differ in abundance between social and solitary bees. Phylogenetic reconstructions based on whole-genome alignments indicate that *Sodalis* has independently colonized halictids at least three times. These strains appear to be mutually exclusive within individual bees, although they are not host-species-specific and no signatures of vertical transmission were observed, suggesting that *Sodalis* strains compete for access to hosts. *De novo* genome assemblies indicate that these three lineages are subject to widespread relaxed selection and that *Sodalis* is undergoing genome degeneration during the colonization of these hosts.

## Introduction

Close contact between individuals of social species provides a means of continual exchange of associated microbes. Indeed, social interactions between hosts substantially impact the composition of bacterial communities associated with those individuals, reducing variation between interactors in honey bees, birds, baboons, and humans (*1*–*5*). Such microbial sharing has even prompted theory suggesting that the advantage of transmitting beneficial microbes could itself select for the evolution of social behavior (*6*, *7*). However, the role that social behavior plays in shaping microbial communities (and perhaps even the reverse) remains an unanswered question.

There are several hypotheses regarding the impact of sociality on gut community evolution. First, because social interactions and cohabitation homogenize bacterial communities among group members (*3*), solitary species should be expected to show more variability in their gut microbes when compared to social relatives. Second, because the establishment of tightly coevolved symbionts requires a dependable mode of intergenerational transmission, we may therefore expect that symbiosis will more readily evolve in social than solitary taxa.

Hymenoptera (bees, ants, and wasps) are an ideal model for studying the feedback between sociality and bacterial communities. Social behavior has evolved a number times in this group, giving rise to the highly eusocial ants and honey bees, but it has also been repeatedly lost in some clades, generating substantial variation in social structure. All Hymenopteran species provision their young with food (*8*, *9*), potentially allowing for some transmission of bacteria from mother to offspring, but the repeated interactions among adult females in social colonies, including trophallaxis in adults in many species (*8*), generates more opportunities for microbial transfer within social taxa.

The vast majority of studies on social insects and their microbiota have examined honey bees and bumble bees, where the microbial communities present appear to be consistent across individuals and generations. The bacterial communities of both of these groups are composed almost entirely of a variety of strains from eight phylotypes spanning five classes (*10*–*14*), several of which have now been implicated in immune and nutritional roles for their hosts (*15*–*17*). In these bee species, strong evidence points to the importance of social behavior in establishing and maintaining these bacterial taxa throughout evolutionary history (*18*) and workers isolated from their nestmates do not host typical microbial communities (*2*, *19*, *20*).

Surprisingly, very few studies have examined differences in the bacterial communities of social and solitary species. Those that have compared social forms have focused primarily on the bee family Halictidae, where social behavior has arisen 2-3 times and has been lost at least a dozen times. Studies of the microbial communities of halictids have compared a handful of social, solitary, and polymorphic species (capable of producing both social and solitary nests within the same population) and have found limited evidence for an effect of social structure on host microbial communities. The first cross-species comparisons suggested that sociality may indeed provide more consistent associations between their hosts and putatively beneficial microbes (*21*), but a subsequent comparison between social and solitary individuals in two socially polymorphic species found no effect of social structure (*22*). It is important to note, however, that in these particular socially polymorphic species, *Megalopta centralis* and *M. genalis*, the bacterial communities do not evolve over long time periods in a strictly social host because females from a single population can produce either social or solitary nests. Therefore, inferences about microbial coevolution with social hosts are limited in this system.

It is unknown whether specialized symbioses originate more readily in social compared to solitary taxa. Obligate symbionts are common across insects and many are only able to survive inside the specialized organs or cells of their hosts (*23*, *24*). The best-studied of these symbionts are the *Buchnera* inhabiting aphid species, many of which are communal or even eusocial (*23*, *25*, *26*), but similar relationships span a diversity of insects including the obligately eusocial ants (*24*, *27*, *28*), flies (*29*–*32*), and weevils (*33*–*35*). Although a number of the taxa identified as important members of the honey bee and bumble bee gut communities appear to be host-specific (*16*, *21*), none have yet been described as obligate inhabitants of bees, suggesting that such relationships may be absent in these insects. However, McFrederick & Rehan (*36*) identified the bacterial 16S rRNA gene sequence of a bacterium in the genus *Sodalis* in the pollen provisions of the bee *Ceratina calcarata*. Nearly every lineage of *Sodalis* previously identified has been found to depend on insect hosts. While the single free-living form is potentially a widespread associate of plants (*37*), its presence in pollen provisions is, nevertheless, quite intriguing.

In order to study the interaction of social behavior with bacterial communities and explore the possibility of bee-associated obligate symbionts, we examined the gut communities across social and solitary species of halictid bees. Because sociality has evolved 2-3 times independently within this clade and has been secondarily lost many more times, halictids have a phylogeny rich in closely related species with different social behaviors (*38*–*42*). We also examined differences in the microbial communities of a single, socially polymorphic species, *Lasioglossum albipes*. Unlike previously studied polymorphic species, the social polymorphism in *L. albipes* has a genetic underpinning (*43*). Variation in social behavior is fixed between well-characterized social and solitary populations, thus providing consistent social or solitary host environments for the establishment and maintenance of microbial communities across generations. Using both these cross-species and intraspecific comparisons, we assess two main hypotheses. First, do the bacterial communities of social hosts differ in composition or variation to those of solitary hosts? And second, is the frequency of incipient symbiosis greater in social hosts than solitary ones?

## Results

## Few taxa dominate the halictid microbiome

We collected 336 samples from western Europe (Fig. 1A) from 23 species of halictids including 11 social species, 5 solitary species, 4 socially polymorphic species, and 1 parasitic species spanning tens of millions of years of evolutionary history (Fig. 1B; Table S1). We sequenced the bacterial 16S rRNA gene using the standard approach of the Earth Microbiome Project, generating 9.9 million read pairs. These sequences were reference clustered into 97% OTUs against the Greengenes database (*44*) using UPARSE (*45*) and QIIME (*46*). Sequences that failed to cluster against this database were subsequently clustered into *de novo* OTUs. Throughout, OTUs with representative sequences in the Greengenes database are referred to using their index number prefaced by “GG” and *de novo* OTUs are prefaced by “denovo”. Details are provided in the Supplementary Information and the full OTU table is provided as Table S2. Rarefaction curves indicate that our specimens were well-sampled (Fig. S1).

**Figure 1.**
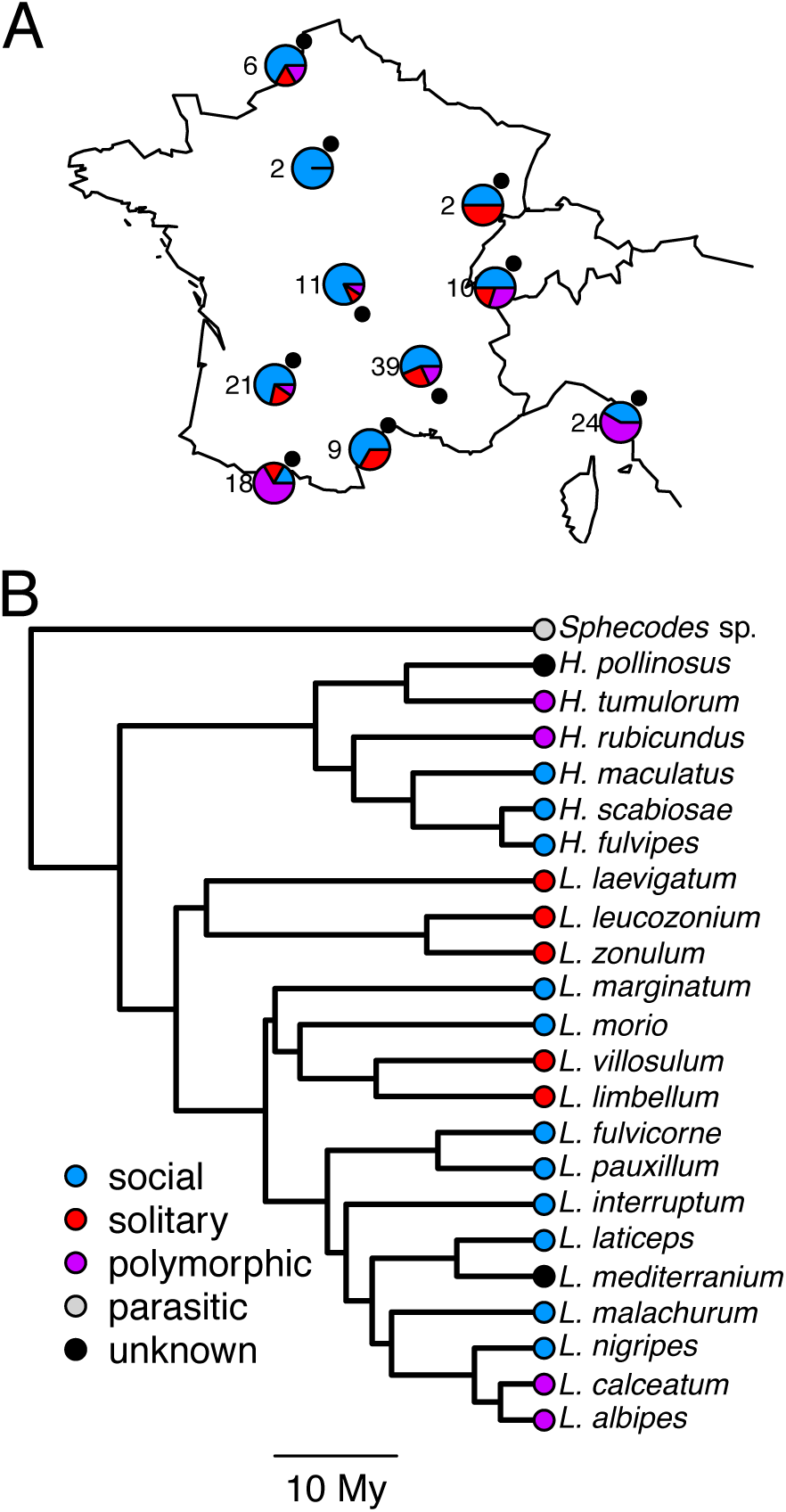
A. Collecting sites for all specimens, number of total specimens from each site, and proportions of those samples classified as each behavioral type. B. Phylogeny of halictid species sampled in this study (adapted from (*41*)). Social behavior is indicated by colors. Branch lengths are based on the time-calibrated phylogeny from (*41*).

The vast majority (97.9%) of sequences recovered from halictids were represented by only the 25 OTUs making up more than 0.1% of total sequences (7,543,271/7,701,949) (Fig. 2A, S2). Among these, three are clear outliers. Greengenes taxon GG273974 (*Wolbachia*), makes up 51.8% of all sequences and taxon GG829017 (*Lactobacillus*) makes up 17.9% of sequences. Lastly, taxon GG4316320 (*Sodalis*) makes up 9.3% of all sequences. All other taxa each make up less than 4% of total sequences.

**Figure 2.**
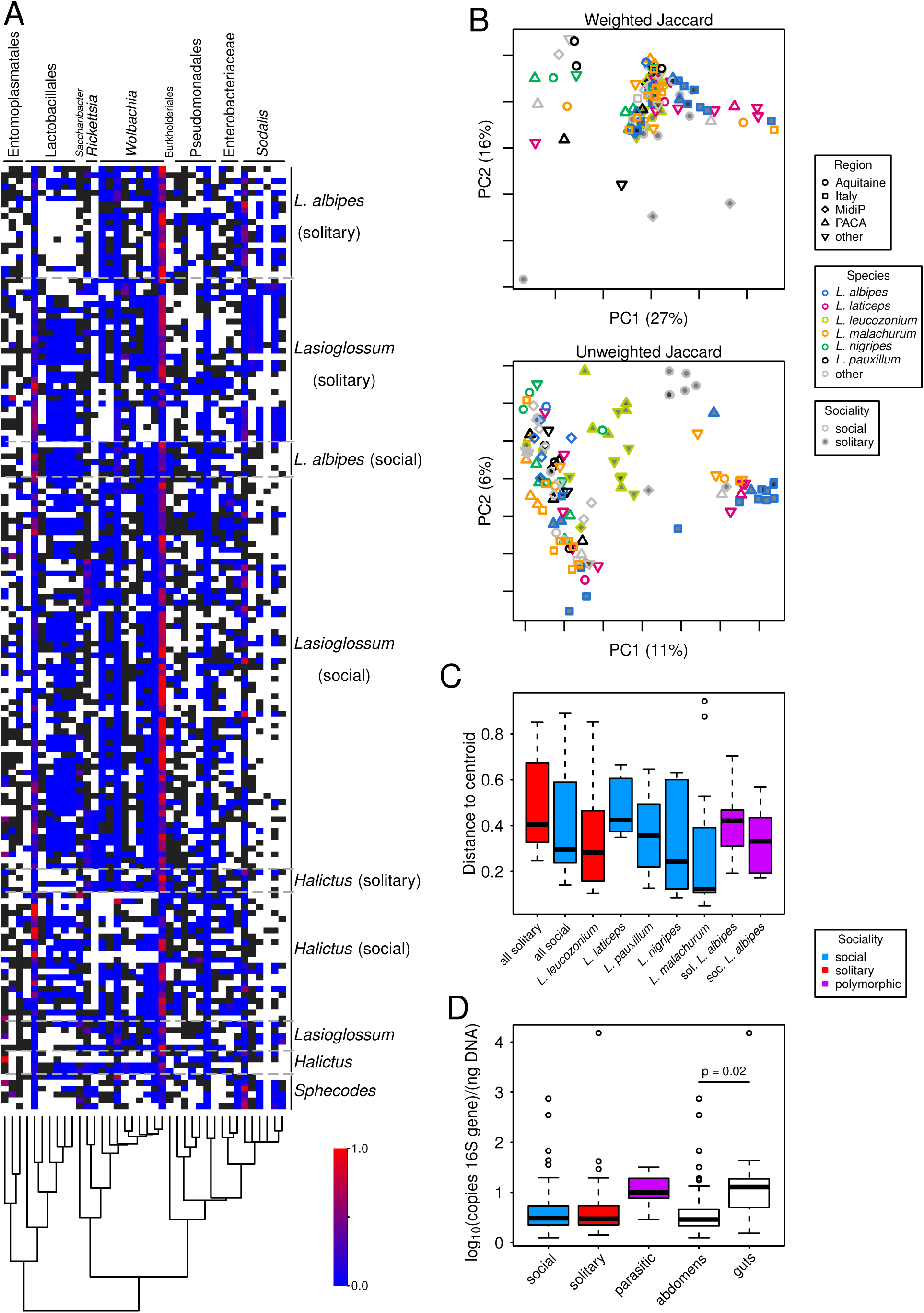
A. Heatmap of relative proportions of the 24 most common bacterial taxa across all samples. When a particular OTU is present in a sample at less than 0.01% frequency, it is colored dark gray. Samples are grouped by genus and social behavior. *Lasioglossum* and *Halictus* samples without a specified behavior are from samples with unknown sociality. Bacterial phylogeny and taxonomy are derived from Greengenes. B. PCoA plots of all halictid samples examined in the current study. “MidiP”: Midi-Pyrénées, “PACA”: Provence-Alpes-Côte d’Azur. The most frequently represented geographic regions and species are indicated and others are grouped together. Individual plots represent the use of different beta diversity metrics (weighted and unweighted Jaccard). C. Distributions of community dispersion in all social and solitary samples as well as those species represented by at least 10 individuals. The polymorphic *L. albipes* is split into social and solitary categories (n=19 solitary, n=6 social). Dispersions are not significantly different between social and solitary bees based on Wilcoxon rank-sum tests. The same test was used to compare all strictly social species to the strictly solitary species *L. leucozonium* and, again, none were significant. Data for *L. albipes* are included for reference; they were not included in statistical comparisons. D. Boxplots showing the number of copies of the bacterial 16S rRNA gene per nanogram of DNA.

Of the 25 abundant OTUs, 16 were also core OTUs (present in at least 50% of individuals). There are an additional 13 core OTUs not represented in the set of abundant OTUs. These 38 core and abundant OTUs include just six *de novo* OTUs not represented in the Greengenes database. Two of these are classified as *Wolbachia* (denovo7 and denovo8), three are classified as Lactobacillales (denovo9, denovo32, denovo63), and one is classified as Enterobacteriaceae (denovo1). The three Lactobacillales are 97% or more similar to a 16S sequence previously recovered from flower nectar (GenBank ID: KX656663). The closest hit in GenBank for denovo1 is the *Sodalis* endosymbiont of the weevil *Sitophilus oryzae* (CP006568) with 93% identity.

We also identified those OTUs present in at least 50% of all samples of *Lasioglossum leucozonium, L. laticeps, L. malachurum, L. pauxillum*, and *L. nigripes*. These species were chosen because they are all represented by at least 10 samples in our dataset. Between 26 (*L. malachurum*) and 69 (*L. laticeps*) core OTUs were identified in each species and between 4% (*L. malachurum* and 59% (*L. nigripes*) of these were not already represented in the set of halictid core OTUs. Taken together, there were 91 additional core OTUs from these species in addition to those identified as core or abundant OTUs in the overall halictid sampling. The resulting 129 OTUs were used for all comparisons of abundance between bee groups.

### Relative abundance of bacteria differs little

We aimed to identify differences in bacterial communities of social and solitary bees. So as not to be confounded by differences between genera and polymorphic taxa, this analysis was limited to only those species in the genus *Lasioglossum* that have been verified as either strictly social or strictly solitary. Mann-Whitney *U* tests of the 38 taxa making up at least 0.1% of all sequences in the dataset or present in at least 50% of all samples show that five 97% OTUs are significantly (FDR-corrected P < 0.01; Table S3) more abundant in solitary bees including one from *Wolbachia* (GG836919), three from *Sodalis* (GG261110, GG2093965, GG4335746), and one from Enterobacteracaeae (denovo1). Four 97% OTUs are significantly more abundant in social bees including three from *Wolbachia* (GG273974, GG835499, GG101940) and one from *Sodalis* (GG4316320). However, as geography may impact the bacteria present, we expanded this analysis to include collection locality as a random effect in a linear mixed model. When geography was taken into account most OTUs were no longer significantly different in relative abundance between social and solitary bees (P > 0.05 using Kenward-Roger approximations for degrees of freedom), with the exception of *Sodalis* OTU denovo1 (t = 2.4, P = 0.046) and *Wolbachia* OTU GG835499 (t = 2.5, P = 0.048), which remained enriched in social and solitary bees, respectively.

Among those additional taxa that were identified as core members of species-level communities but were not core taxa among all halictids, 15 were significantly more abundant in solitary samples (FDR-corrected P < 0.01; Table S3). These include nine OTUs classified as Enterobacteriaceae, five OTUs classified as *Sodalis,* one classified as *Erwinia,* and one classified as *Wolbachia*, many of which were completely absent from social bees (Table S3). No OTUs were significantly more abundant in social samples. Unfortunately, the small number of samples in which many of these taxa were present prevented the inclusion of geographic region in the model.

We also implemented Mann-Whitney *U* tests between social forms of the polymorphic species *L. albipes*. A single OTU (GG830148: Lactobacillales) was significantly more abundant in social samples after FDR correction (P < 0.01). No taxa were significantly more abundant in solitary samples. Unfortunately, we are unable to control for geographical differences in *L. albipes* because differences in social behavior occur across populations in this species, and thus covary with geography.

We also tested for differences in the frequency of *Sodalis* within social forms of *L. albipes* through the detection of *Sodalis* reads in an *L. albipes* shotgun genomic resequencing dataset. Of 75 specimens of each behavioral type, only six social samples have detectable levels of *Sodalis* as opposed to 30 solitary samples (Supplementary Information). This difference in frequency is significant according to a X^2^ test (P = 1.1x10^−5^, X = 19.3). Again, sociality covaries with geographic region in this species so these results may be confounded by geographic variation. However, it is worth noting that *Sodalis* is present in other halictid species from all three of the geographic regions from which these social samples were collected (Rimont, Dordogne, and Calais; Fig. S2) indicating that geography is unlikely to be the factor limiting *Sodalis* colonization.

We also used Mann-Whitney *U* tests of the copies of bacterial 16S rRNA gene per nanogram of DNA to test for differences in total quantity of bacteria in social and solitary samples. Quantities did not differ between social and solitary bees (P > 0.1) or between either social or solitary bees and parasitic bees. In general, the abundance of bacteria does not show any obvious pattern with bee behavior or evolutionary history (Figs. 2D, S3). As expected, we recover a higher relative concentration of bacterial DNA in guts than whole abdomens (P = 0.016).

### Sociality does not affect community variability

We tested for differences in variability of individual bacterial taxa using Brown-Forsythe tests of the 129 core and most common OTUs and found that 2 taxa had significantly different variance in social and solitary bees (FDR-corrected P < 0.01; Table S3). Both of these taxa had higher variance in social samples.

We also compared variance of community assemblage by examining dispersion of weighted (F = 3.2, P = 0.076) and unweighted Jaccard distances (F = 0.38, P = 0.54), but did not find significant differences (Fig. 2C). We also compared the solitary *L. leucozonium* to all social taxa with at least 10 samples using both metrics but only a single comparison showed any indication of a difference in community dispersion. *L. laticeps*, a social species, had greater community dispersion using weighted Jaccard (F = 4.41, P = 0.047).

### *Sodalis* and *Wolbachia* dominate community differences among social forms

No clear differences in overall bacterial communities of bees with different behaviors, from different species, or collected at different locations are apparent from PCoA analyses (Fig. 2B). We used automated supervised learning classification to determine whether these communities were distinguishable in any way. For the classifier of samples based on social behavior, the ratio of baseline to observed error was 2.80 meaning that the classifier was 2.8 times better than random guessing. We consider error ratios ≥ 2 to indicate that there are significant differences in communities (*47*, *48*). In this classifier, all social samples were correctly classified and 18/28 (64%) solitary samples were correctly classified. Of the 10 most important features for distinguishing social behaviors, nine are classified as *Sodalis* or the family to which it belongs, Enterobacteriaceae, including three of the most abundant OTUs (GG261110, GG4335746, denovo1). The last OTU in the top 10 is from *Wolbachia* (GG836919). In order to determine whether other taxa had similar discriminatory power, we removed all 451 OTUs classified as Enterobacteriaceae and reran the supervised learning, which reduced the error ratio to 1.28 with 56/57 social samples correctly classified but only 7/28 (25%) solitary samples correctly classified.

From this analysis, we can conclude that *Sodalis* is by far the most important clade distinguishing social and solitary taxa. Indeed, building a classifier based only on the 42 *Sodalis* OTUs is more accurate than one based on all data, with an error ratio of 3.58 and 57/57 social samples and 20/28 (71%) solitary samples correctly classified. Frequency differences of *Sodalis* are apparent between these two groups (Fig. S4).

The results when classifying communities from five bee species represented by at least 10 samples in our dataset had an error ratio of 2.71. However, removing the 127 *Wolbachia* and *Rickettsia* OTUs decreased the error ratio to 1.57. A classifier based entirely on the 127 *Wolbachia* and *Rickettsia* OTUs was quite successful with an error ratio of 2.42. Therefore, at least among the five bee species included in this analysis, there is significant species-specificity of *Rickettsia* and, to a lesser extent, *Wolbachia.* While few recovered OTUs are unique to particular species, there are clear differences in frequency of individual taxa (Fig. S5). Results of all supervised learning analyses conducted are presented in Table S4.

### Alpha diversity

We do find that, when including all *Lasioglossum* samples, OTU counts are greater in solitary than social samples (Wilcoxon rank-sum test P = 0.01). The Chao1 index also supports this trend (P = 0.04). We, therefore, tested for differences in alpha diversity between individual pairs of social and solitary bee species with at least 10 specimens as including multiple species in a group is likely to impact alpha diversity measures. All tests between the solitary *L. leucozonium* and social taxa (*L. malachurum, L. laticeps, L. pauxillum,* and *L. nigripes*) are not significant (P > 0.05) except for *L. malachurum* (P = 7.9x10^−5^) for OTU counts and for Chao1 (P = 0.0005).

We also tested for differences in variance of alpha diversity in social and solitary samples using Brown-Forsythe tests. When including all *Lasioglossum* specimens, both OTU counts (F = 4.7, P= 0.04) and Chao1 (F = 4.7, P = 0.04) appear to have greater variance in solitary samples. However, when testing between individual species, these tests are not significant (P > 0.05) except between *L. leucozonium* and *L. malachurum* for both OTU counts (F = 22, P = 5.66x10^−5^) and Chao1 (F = 15.3, P = 0.0005). Evidence for greater variance in alpha diversity of solitary bees is, therefore, extremely weak.

### *Sodalis* phylogeny

The presence of *Sodalis* was unexpected, particularly at the high abundance seen. We therefore mined the *Sodalis* data from several previously collected genomic sequencing datasets, including 150 samples of the socially polymorphic *Lasioglossum albipes* and individual samples of 12 other halictid species. We also included all shotgun sequencing data collected for the five bee genomes sequenced in Kapheim et al. (*49*) and *Ceratina calcarata* (*50*). We recovered large amounts of *Sodalis* data from 36 of the 150 *L. albipes* samples as well as six other halictids (*H. ligatus, L. calceatum, L. leucozonium, L. malachurum, L. marginatum,* and *L. vierecki*) and *C. calcarata* (Fig. 3). Four lineages of bee-associated *Sodalis* are apparent from this inference. The lineage inhabiting *C. calcarata* was distantly related to all others, closer to the *Sodalis* symbiont of tsetse flies than to any halictid-associated *Sodalis*. The *L. leucozonium* lineage was also unique among the samples we examined and we named this *Sodalis* lineage “SLEU”. The rest of the samples fell into two large clades. The first (“SAL1”) includes 12 *L. albipes* samples as well as all of the other halictid samples. The last halictid-associated clade (“SAL2”) is composed entirely of *L. albipes* samples. Based on these results, we used metagenomic approaches to assemble genomes *de novo* for SLEU, SAL1, and SAL2 (Supplementary Information). We also attempted to assemble the genome of the taxon associated with *C. calcarata* but were unsuccessful.

**Figure 3.**
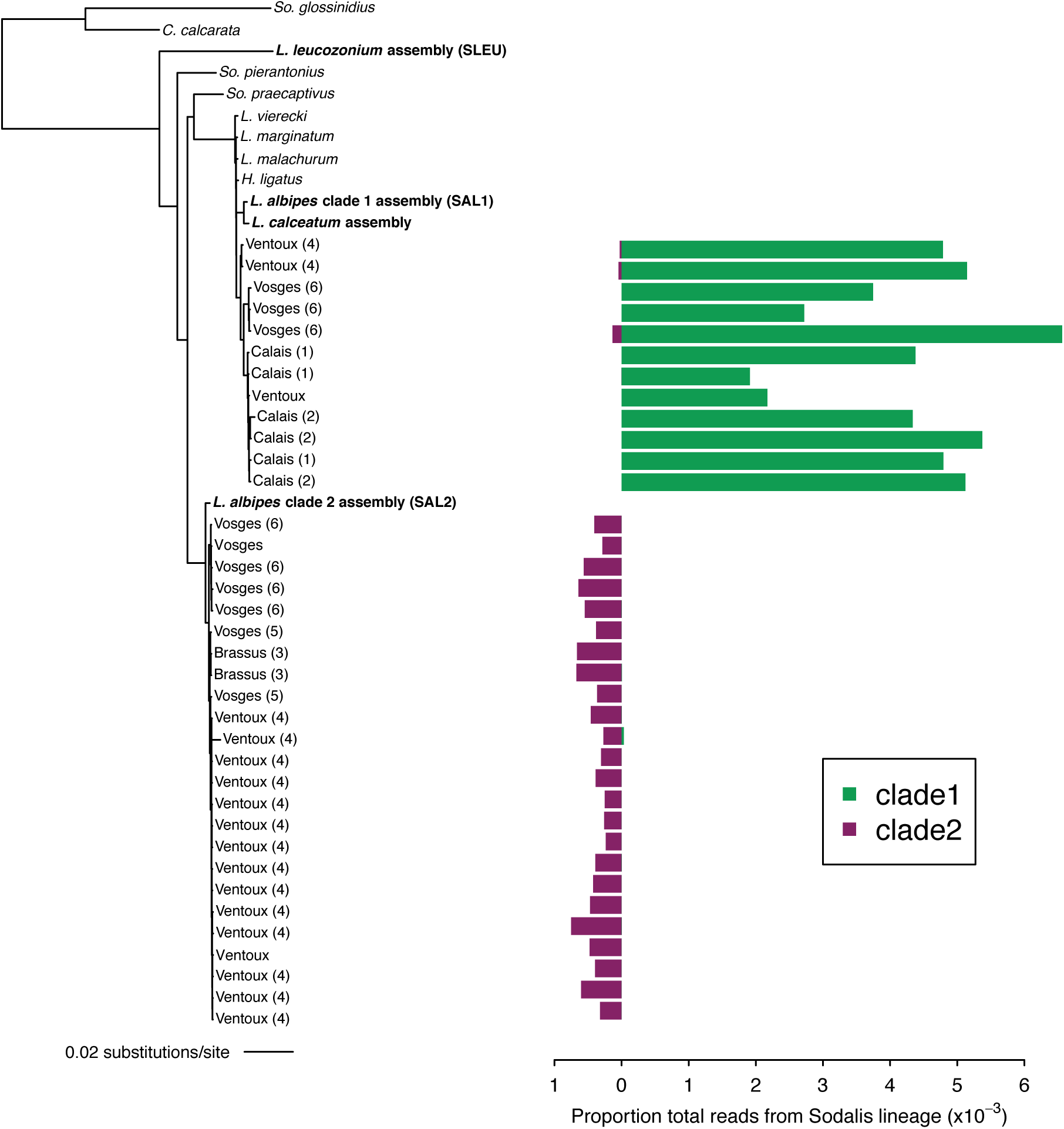
Phylogeny of recovered *Sodalis* sequences. Bolded taxa represent genomes assembled *de novo* in this study. *C. calcarata, L. vierecki*, *L. marginatum*, *L. malachurum*, and *H. ligatus* all represent *Sodalis* sequences recovered from shotgun sequencing libraries derived from those bee taxa. *So. glossinidius* and *So. pierantonius* are previously assembled endosymbionts of tsetse flies and weevils, respectively. *So. praecaptivus* is the previously assembled free-living form of *Sodalis* (*33*). All other tips are derived from *L. albipes* shotgun sequencing libraries and are labeled with their location of origin and mitochondrial haplotype in parentheses. Each number represents a unique haplotype. We were not able to recover haplotypes for all samples. Bars represent the proportions of total reads that derived from the two *Sodalis* lineages found in *L. albipes* within each *L. albipes* sample.

The two clades of *Sodalis* that include the *L. albipes* samples do not show any apparent segregation by collecting locality (Fig. 3). Two of four localities are shared between the two lineages suggesting that both lineages are widespread and potentially interact within the same bee populations. It also appears that all three *Sodalis* lineages occupying halictids have convergently evolved this lifestyle. SLEU in particular is quite distantly related to the other clades occupying halictids.

### *Sodalis* genome characteristics

Separately assembling data from each identified clade of *Sodalis* yielded unique genome sequences. SAL1 was composed of 20 scaffolds totaling 4.1 Mb (though 527 kb are composed of gaps) and the largest scaffold making up 3.2 Mb of that assembly. RAST annotated 4,314 genes in this genome and it has an average GC-content of 56.7%. SAL2 has a similar total length of 4.2 Mb but these are placed in 241 scaffolds, the largest of which is 83 kb with an N50 size of 27 kb and a GC-content of 57.5%. Only 69 sites in this assembly are uncalled. There are 4,518 genes in this assembly. Lastly, the SLEU assembly is much smaller with a total length of 2.4 Mb on 86 scaffolds and the largest scaffold is 530 kb with an N50 of 372 kb, though 878 kb are gaps. RAST identified 2,404 genes in this genome and it has a %GC of 54.4. By comparison the free-living form of *Sodalis*, *S. praecaptivus*, has a GC-content of 57.5%, a length of 4.7 Mb and 4,411 genes annotated by RAST. Particularly for SLEU, the decreased length and GC-content in the halictid inhabitants both suggest that these genomes may be degenerating (Supplementary Information).

All three genomes were relatively complete. SAL2 has 98.8% of the expected 137 marker genes, SAL1 has 98.2%, and SLEU has 92.8%. Interestingly, coding density was also found to vary widely. Over the whole sequence, 74% of the SAL1 genome is composed of coding sequence, 84% of SAL2 is coding sequence, and only 58% of SLEU is coding sequence.

### Coinfections are rare

The finding that two *Sodalis* lineages coexisted in *L. albipes* even within the same geographic populations suggested that these lineages may interact directly. We examined the degree to which these two lineages occurred together by analyzing diagnostic SNP’s for each clade and identified 6,407 such sites which allowed for confidence in distinguishing these two groups (Supplementary Information). Although we detect reads from both strains in all but two bees, one strain always dominates and there are only two individuals in which each strain makes up at least 1% of all *Sodalis* reads. The frequency of these coinfections with appreciable numbers of reads from both strains are less common than expected by chance (Fisher’s exact test P = 0.044; Fig. 3). This pattern could be explained by competition between the two lineages or by vertical transmission of *Sodalis* from mother to offspring.

### No evidence for long-term vertical transmission

Many obligate endosymbionts are transmitted vertically from mother to offspring, serving to enforce strict coevolution between the host and symbiont. In order to determine whether this mode of transmission is dominant in *Sodalis*, we compared the presence of *Sodalis* lineages in the *L. albipes* samples with the maternally inherited mitochondrial haplotypes of those same samples. If vertical inheritance is the primary mode of transmission, then mitochondrial haplotypes may co-occur with a single lineage of *Sodalis.* However, we found no indication of long-term maternal inheritance (Fig. 3).

### Widespread relaxed selection in *Sodalis*

Purifying selection is often relaxed in obligate endosymbionts as large parts of their own genomes are no longer necessary for their survival and as the result of population bottlenecks during colonization. Within *Sodalis*, relaxed selection has been previously identified in both *S. pierantonius* and *S. glossinidius* (*33*, *51*). We inferred the presence of relaxed selection by examining genome-wide dN/dS distributions. All *Sodalis* endosymbiont genomes presented in this study as well as the weevil endosymbiont *S. pierantonius* have significantly higher distributions of dN/dS ratios than the only known free-living form of *Sodalis, S. praecaptivus* (P< 2.2x10^−16^; Fig. 4) indicating widespread relaxed selection. Notably, the tsetse endosymbiont, *S. glossinidius* does not have significantly higher genome-wide dN/dS but instead has significantly lower dN/dS (P = 4.2x10^−11^). This may be due to the fact that *S. glossinidius* has occupied tsetse flies for a much longer period of time and the genes that experienced relaxed selection may have been completely lost.

**Figure 4.**
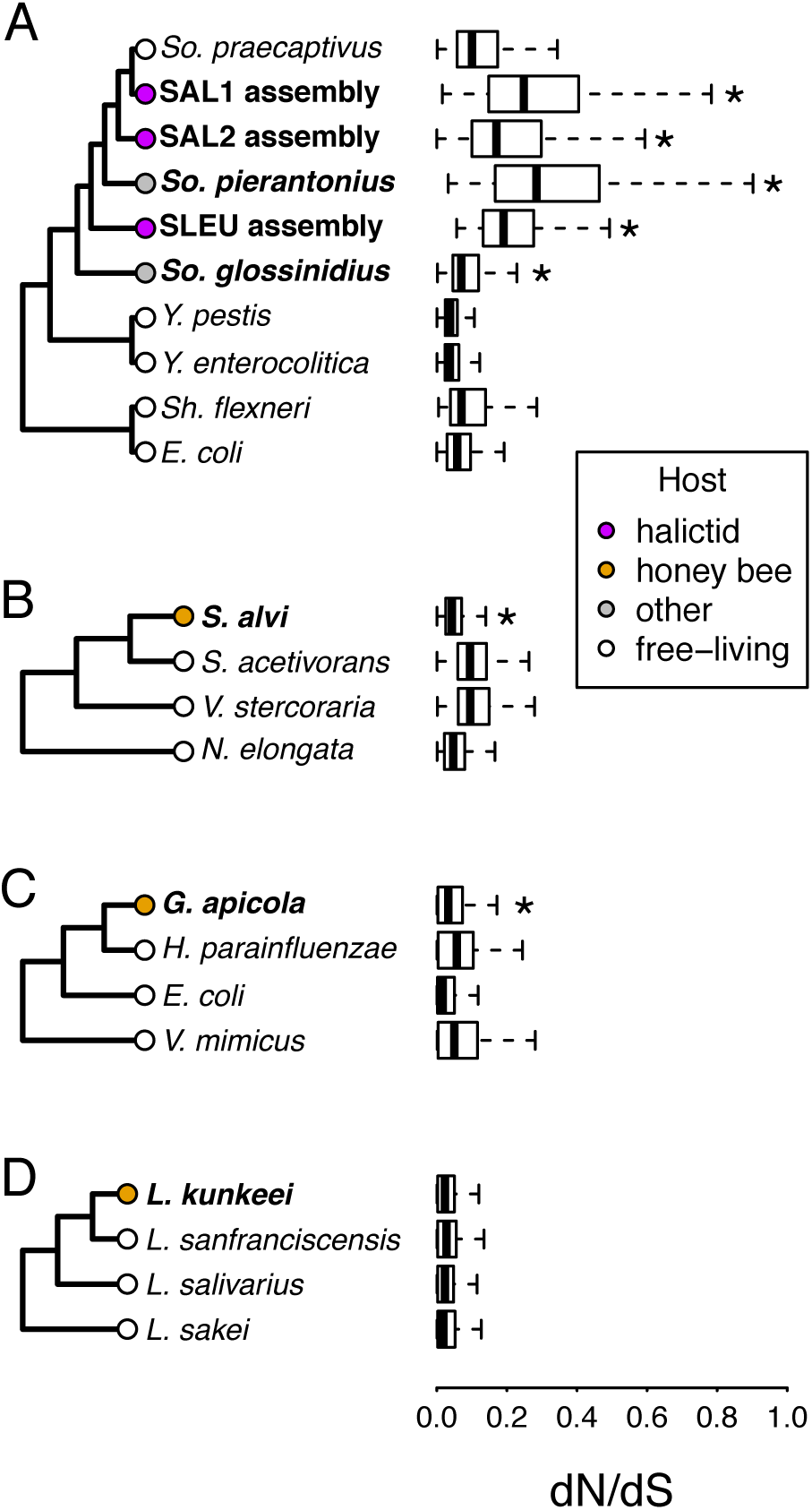
Distribution of dN/dS ratios in the full genomes of four bacterial taxa of interest and their close relatives: A. *Sodalis* endosymbionts including those identified in this study as endosymbionts of halictids, B. *Snodgrassella alvi*, C. *Gilliamella apicola*, and D. *Lactobacillus kunkeei*. These last three taxa are well-known endosymbionts of the honey bee, *Apis mellifera*. Significant differences were determined by comparing each endosymbiotic taxon to the most closely related free-living taxon (e.g. *So. pierantonius* compared to *So. praecaptivus*) using Wilcoxon signed-rank tests.

We also sought to determine whether similar patterns exist for any of the endosymbionts previously identified in honey bees. Both *Snograssella alvi* and *Gilliamella apicola* have significantly lower dN/dS ratios than the most closely related species included in the analysis (*S. acetivorans* and *Haemophilus parainfluenzae*, respectively) and *L. kunkeei* does not differ significantly from its relatives. The genome-wide relaxed selection that the halictid-occupying *Sodalis* lineages are subject to appears to be unique among bees.

### Genes lost across functional categories

We attempted to infer likely functional consequences of *Sodalis* in halictids by examining the numbers of genes assigned to different functional categories. The numbers of genes in each functional subsystem was almost always lower in each of the halictid-inhabiting *Sodalis* taxa, particularly for SAL1 and SLEU, the two lineages with the highest average dN/dS ratios, smallest total sizes, and lowest GC-content (Fig. S4). No clear function of *Sodalis* within their hosts was apparent from this analysis.

### *Sodalis* is present throughout the body

The presence of endosymbionts in particular organs can also inform its functional consequences. We conducted diagnostic PCR’s for the presence of *Sodalis* in three *L. albipes* abdomens, thoraces, heads, legs, and antennae (Supplementary Information). These yielded a product for all body parts for all three samples, except for heads. The amplification of bee COI sequence was also unsuccessful for heads indicating that this DNA extract was likely of too low quality for PCR. While their abundances likely vary in different body parts, these results suggest that *Sodalis* are found throughout the body and are likely present within the hemolymph of bees, as seen for *S. glossinidius,* the *Sodalis* symbiont of tsetse flies (*52*, *53*) and a number of other vertically transmitted, non-gut symbionts.

## Discussion

Our results suggest that social behavior has a limited influence on the microbial communities of social and solitary bees. Our cross-species comparisons revealed a consistent association of social structure with the prevalence of only one bacterial taxon: *Sodalis.* We also compared social and solitary females in a socially polymorphic bee species, *L. albipes*. Even within this polymorphic species, only Lactobacillales and *Sodalis* showed any signature of variation across social forms.

Given that research has clearly shown the importance of social interactions in the establishment of bacterial communities in honey bees (*2*), it is surprising that we find so few differences in communities of social and solitary halictids. However, our findings are consistent with previous halictid studies, where comparisons of two sympatric and socially polymorphic species found no effect of social structure on host bacterial communities (*22*). However, in those species, the polymorphism occurs among females within the same population, suggesting that the symbionts would not evolve under consistent host social conditions. In contrast, the social polymorphism in *L. albipes* has a genetic underpinning and the differences occur across populations (*43*), thus generating a consistent host environment.

Our expectation that microbes would be more readily shared in social bees is based largely on our assumptions about the frequency of trophallaxis and physical contact between nestmates. Many social (but not solitary) halictids have been observed repeatedly opening, cleaning, and inspecting the cells of their developing offspring (*8*, *54*), and it is abundantly clear that there is a greater extent of interactions among adults in social versus solitary halictid nests. However, both social and solitary halictids are mass provisioners, collecting and provisioning the full pollen supplies to their developing offspring at the time of oviposition (*8*) and, in addition, bacteria can easily colonize host bees from the environment (*21*, *55*–*57*). These latter modes of transmission now appear to be much more influential than we initially predicted.

It is notable that those taxa that appear to be most dependent on social interactions for their stable establishment in honey bees (*Gilliamella apicola*, *Snodgrassella alvi,* and *Frischella perrara* (*2*)) are also those for which an equivalent symbiont is not apparent in halictids. On the other hand, *Lactobacillus* establishes in quantity even when newly emerged honey bees have only limited exposure to nest materials (*2*), is reliably found in many bees and ants (*58*, *59*), and is the second-most abundant taxon found in halictids. These findings all serve to show that *Lactobacillus* robustly colonizes Hymenopteran guts and may be among the most universal members of bee-associated bacterial communities.

Only *Sodalis, Wolbachia,* and *Rickettsia* show any consistent differences between halictid species or behavioral types. Given that *Wolbachia* and *Rickettsia* are intracellular and vertically transmitted, their specificity is unremarkable and has been shown previously (*60*). Far less is known about *Sodalis* in bees and, though it has been previously identified in association with the bee genera *Ceratina* and *Osmia*, (*36*, *61*), it has not previously been examined in detail. The diversity of insects that play host to *Sodalis* is quite wide, spanning lice, beetles, flies, many hemipterans, and, now, bees (*31*, *51*, *62*–*67*). Where best-studied, in the rice weevil *Sitophilus oryzae* and the tsetse fly *Glossina morsitans*, the functional consequences of *Sodalis* symbiosis appear to vary. The weevil symbiont, *Sodalis pierantonius,* plays an essential role in exoskeleton deposition in its hosts and is eliminated from all organs except the ovaries after the exoskeleton has been fully formed (*35*). However, no clear function is apparent for *S. glossinidius* in tsetse flies, where the symbionts show comparatively less regular localization (i.e. found in hemolymphy) than seen for weevils (confined to bacteriocytes) (*52*, *53*). The location of *Sodalis* in its Pthirapteran hosts appears to be tissue-specific (*62*) and varies from bacteriocyte occupation in spittlebugs (Hemiptera: Auchenorrhyncha: Cercopoidea) (*66*) to *S. glossinidius*-like occupation of various tissues in stinkbugs (*67*). Our ability to amplify *Sodalis* from both the legs and antennae of bees suggests that it may exist as a widespread hemolymph endosymbiont.

Given the number of different insects occupied by *Sodalis* and the diversity of its consequences in these taxa, it is premature to draw any conclusions about its putative function in bees. However, our ability to distinguish social and solitary halictids based entirely on the *Sodalis* strains present combined with the weak but repeatable evidence for differences in *Sodalis* abundance between the two behavioral types is suggestive. This pattern is not the result of phylogenetic relationships; these differences occur over multiple independent lineages that have convergently lost social behavior. It is possible that “social immunity”, the group-level behaviors that social species can use to combat pathogens (*68*), may form the basis of these differences. The significantly more frequent occurrence of *Sodalis* in solitary samples both across halictids and within the socially polymorphic species *L. albipes* is consistent with *Sodalis* being more readily eliminated or prevented from establishing in social bees.

Our study shows that *Sodalis* is predisposed to evolving symbiosis with insects. Including the halictids in this study, at least 12 evolutionarily distinct groups of insects have been separately colonized by *Sodalis*. Several of these lineages show evidence for relaxed selection and rapid genome degeneration, implying nascent obligate symbiosis (*33*, *51*, *63*, *69*, *70*). Even in comparison to *Wolbachia*, which is estimated to occupy around 20% of insects (*71*), the flexibility of *Sodalis* is impressive. We find that at least three strains of *Sodalis* have independently colonized a single genus of halictid and a fourth strain, apparently spawned from the same ancestral lineage as *S. glossinidius,* occupies *C. calcarata,* a species more closely related to honey bees than to halictids. Yet we find little evidence for host specificity or, given the presence of the same strain in both European and American bees, geographic limitation of individual strains. Even within a single species of halictid (*L. albipes*) we clearly identified two distinct *Sodalis* lineages that have independently colonized this species, showing a remarkable predisposition of free-living *Sodalis* to infect halictids. And most excitingly, these two *Sodalis* lineages exist in a mutually exclusive way, suggesting that they may be competing for access to host space. The widespread relaxed selection in the genomes of these *Sodalis* lineages shows that they are also undergoing genome degeneration as has been found previously. Yet here we have potentially added another fold to our understanding of symbiont biology: competition between distinct evolutionary lineages of a nascent endosymbiont for full access to hosts.

Werren & Windsor (*71*) proposed that *Wolbachia*’s prevalence indicates a global equilibrium or, alternatively, an ongoing increase in the frequency of *Wolbachia* infection. We propose that a third model may be more appropriate for *Sodalis*. In addition to the four strains identified in bees, the free-living form of *Sodalis* appears to have independently colonized each of the other insect groups with which it associates (*33*, *37*, *72*, *73*). The genomes of each of the lineages that have been sequenced all show widespread relaxed selection, implying that, in every case, *Sodalis* is in the process of becoming an obligate endosymbiont. However, the frequency with which *Sodalis* infections are established suggests that it may never reach stability as an obligate symbiont but each lineage may, instead, be eliminated from a given host, either by host immune function, random events, or competition with a more recently colonizing strain. Rather than progressing towards stable symbiosis, this may be a continually renewing process wherein new strains are constantly established and eliminated.

Given the differences between honey bee and halictid-associated bacteria, and the relative consistency of these communities regardless of host behaviors, sociality clearly does not lead to a particular bacterial community. However, we do find evidence that social behavior influences the abundance of several strains of *Sodalis* both within a single socially polymorphic species and between strictly social and solitary taxa. The mechanism of this impact is, apparently, specific to the conditions of the interactions between *Sodalis* and their hosts as this pattern is unique among the bacteria examined. However, contrary to our prediction that symbiosis would occur more readily among social hosts, the prevalence of *Sodalis* is higher in solitary bees, tentatively suggesting that social species may be more effective at purging these strains than solitary ones. Unfortunately, until a clear determination can be made as to the nature of the interaction between *Sodalis* and halictids and why abundances differ between bees with different behaviors, we cannot draw any conclusions about whether social evolution is correlated with the presence of beneficial microbes (*6*, *7*). The identification of an apparent incipient symbiont in this socially variable clade of bees does, however, provide a compelling system for understanding the possible role of host social behavior in this process.

## Methods

### Sample collection

Bees were collected from 10 regions in France, Italy, and Switzerland (Figs. 1A, S2) between 2010 and 2014 and were immediately flash frozen over liquid nitrogen. Characteristics of all samples are given in Table S1. For most samples, we used whole abdomens for DNA extraction, but guts were also dissected under sterile conditions for 30 specimens. We used the Mo Bio PowerSoil DNA Isolation Kit (Qiagen, Hilden, Germany) with the addition of proteinase K digestion (*74*) for all DNA extractions.

### Quantifying absolute abundance of bacteria

Triplicate quantitative-PCRs were performed as in Rubin et al. (*74*) and Sanders et al. (*75*). We took the average of the triplicate measurements for each specimen and standardized these measurements to the concentration of overall DNA present in each sample, measured using a Qubit Fluorometer (Thermo Fischer Scientific, Waltham, MA, USA). Numbers of copies of the bacteria 16S gene per nanogram of DNA was compared between social and solitary specimens using Wilcoxon rank-sum tests.

### 16S sequence processing

Bacterial 16S rRNA gene amplicon sequences were demultiplexed with QIIME (*46*) and then processed with the UPARSE (*45*) pipeline to remove low quality and chimeric sequences. The standard operating procedure was followed. Briefly, paired reads were merged, discarding contigs with lengths over 300 bp (median, lower quartile, and the upper quartile were all of length 253) or more than 0.5 expected errors. Singleton sequences were also discarded. Chimera detection was done both with the *de novo* chimera detector built into UPARSE and the UCHIME pipeline using the Greengenes gold database. All filtered sequences were then clustered against these representatives at 97% similarity. Those that did not cluster were discarded.

To compare with other bee microbiomes, we combined our data with two previously reported datasets. Firstly, we drew sequence data from an unpublished Earth Microbiome Project dataset on the honey bee microbiome from Qiita (Study ID: 1064 and in the European Nucleotide Archive under study ID: PRJEB14927). This dataset consisted of 6,744,896 150 base single-end rather than paired-end reads but was otherwise quality filtered using the same procedure as our halictid dataset. We also included the 4,999 bacterial 16S sequences from Martinson et al. (*11*) from GenBank, truncating them to 1,000 bp before clustering.

This combined dataset was open reference clustered against the August 2013 release of the Greengenes database using usearch61 as implemented in QIIME with chimera detection deactivated. Taxonomy for *de novo* OTUs was assigned using RDP (*76*), sequences were aligned against the Greengenes database using PyNAST (*77*) and inserted into the reference Greengenes (*44*) tree using ParsInsert (https://sourceforge.net/projects/parsinsert/). OTUs classified as chloroplast or mitochondria (368) or that failed to align to the Greengenes database (661) were discarded, as were those not represented by at least two sequences (1,663). Halictid samples not represented by at least 20,000 of the remaining reads were also discarded. The OTU table used for all analyses is included as Table S2. The final OTU table includes 7,053 OTUs. After filtering to just halictid samples, 2,039 OTUs remain.

### Behavioral classification

Behavioral information for the bee taxa examined here was drawn from (*41*) and references therein. However, we treated *L. fulvicorne* as social rather than solitary (*54*). Behaviors of the species *L. mediterrraneum* and *L. pollinosus* have not been previously well-characterized, so they were not included in comparisons of social behaviors.

### Differences in social and solitary bacterial communities

After filtering to include only *Lasioglossum* samples with known social behavior, excluding all samples from the behaviorally polymorphic species *L. albipes* and *L. calceatum*, and rarefying the resulting table to 20,000 observations, we tested for differences in the abundance of taxa between social and solitary samples using Mann-Whitney *U* tests. We tested only the core OTUs present in at least 50% of samples overall as well as those present in at least 50% of samples in each species represented by at least 10 samples. We also tested all taxa making up at least 0.1% of sequences in the overall dataset. P-values were corrected for multiple testing using the Benjamini-Hochberg procedure. We also implemented Mann-Whitney tests between social forms of *L. albipes* to identify overlapping differences between and within species. In order to confirm these results, we also performed ANOVA tests on those taxa identified as significantly different in abundance, including geographic region from which they were collected as a random variable.

We hypothesized that solitary communities may be more variable than social communities. We therefore used Brown-Forsythe tests to assess differences in the variance of the core and common taxa in social and solitary bees. We also assessed differences in community dispersion using the PERMDISP2 procedure implemented as betadisper in the R vegan package. We calculated beta diversity for this analysis using both unweighted and weighted Jaccard.

### Classifying by behavior, location, and species

We built automated classifiers using the supervised learning method implemented in QIIME to determine whether bacterial communities were identifiably different between bee species, social behaviors and collecting locality. In this approach, OTUs of samples are used to construct models that predict sample groupings based on those OTUs (*47*, *48*). Error was estimated using 10-fold cross validation, splitting the data into 10 sets with approximately equal group membership and using 9/10 sets to build the classifier and predict membership in the last set. Model accuracy is assessed using the estimated generalization error: the ratio of the proportion of classification errors that would occur if guesses were completely random to the proportion of errors from the model. Higher error ratios mean that the classifier is performing better. An error ratio of ≥ 2 is typically used to indicate that there are significant differences in communities (*47*, *48*). An error ratio of 1 indicates that a classifier is no better than random and that OTUs do not differ predictably between groups. All analyses were conducted on tables with OTU quantities represented as proportions so that data were not lost during rarefaction.

For classification of social behaviors, we used only *Lasioglossum* species with known records of social or solitary behavior, excluding the polymorphic *L. albipes* and *L. calceatum.* We used the same dataset for classification of species with the additional constraint that all species included had to be represented by at least 10 samples. This limited the dataset to representatives of *L. laticeps, L. leucozonium, L. malachurum, L. nigripes,* and *L. pauxillum*. We also built a classifier for collection locality, again limiting the dataset to those regions represented by at least 10 samples (i.e. Aquitaine, Florence, Midi-Pyrénées, and Provence-Alpes-Côte d'Azur). Lastly, we used all *Lasioglossum* and *Halictus* data to classify genera.

### Alpha diversity

We calculated alpha diversity as observed counts of OTUs and the Chao index calculated on 100 replicate OTU tables rarefied to 20,000 reads and tested for differences between social and solitary *Lasioglossum* samples using Wilcoxon rank-sum tests, again excluding the polymorphic and uncharacterized species. We calculated rarefaction curves using mothur version 1.36 (*78*).

### Evolutionary history of halictid-associated *Sodalis*

*Sodalis* was an unexpected yet dominant member of the recovered bacterial communities, so we integrated a number of genomic datasets in order to examine it in greater detail. First, we constructed a phylogeny of the *Sodalis* sequences found in halictids using a whole-genome approach leveraging a previously collected Illumina HiSeq2000 shotgun sequencing dataset of 150 adult male *L. albipes*. Reads that failed to map to the *L. albipes* genome (after it had been filtered for bacterial scaffolds) with BWA (*79*) were used to create a metagenomic dataset with a high-proportion of bacterial reads (Supplementary Information). In addition, we made use of previously collected sequencing data for 13 other halictid species: *Augochlorella aurata*, *Augochlora pura, Agapostemon virescens, Lasioglossum leucozonium, L. figueresi, L. marginatum, L. vierecki, L. zephyrum, L. calceatum, L. malachurum, L. oenotherae, L. pauxillum,* and *Halictus ligatus*. Finally, to determine how widespread *Sodalis* is across bees, we also downloaded all raw data (excluding mate-pair libraries) sequenced as part of the 10 bee genome project (*49*) and for the *Ceratina calcarata* genome sequencing project (*50*). All raw datasets were filtered using the same methodology as the *L. albipes* dataset.

We then downloaded seven Enterobacteriaceae genomes and all coding sequences from GenBank including three *Sodalis* taxa. These genomes were aligned using Mauve (*80*). Initial analyses suggested that *S. pierantonius* is closely related to the halictid-associated *Sodalis*. Therefore, we mapped our sets of reads that failed to map to bee genomes to the genome-aligned *S. pierantonius* using BWA (*79*) and called genotypes using FreeBayes (*81*). The resulting aligned sequences were then used to create a maximum likelihood phylogeny using RAxML v7.3.0 (*82*) specifying a GTRGAMMA model of nucleotide substitution.

### *Sodalis* genome assembly

The recovered phylogeny showed the existence of two distinct clades of *Sodalis* within *L. albipes.* In order to obtain the best possible genome sequences from each of these clades, we assembled the unmapped reads from the individual samples with the highest number of reads mapped to *S. pierantonius* using a metagenomic approach (Supplementary Information). After determining that the lineages inhabiting *L. leucozonium* may be unique and that additional information may be available in the 10x data from the other species, we also assembled *Sodalis* from *L. leucozonium* and *L. calceatum de novo*. Genome completeness for all assemblies was determined using CheckM (*83*).

### Gene tree inference

We endeavored to confirm that our phylogeny was indeed the most likely by examining gene trees. We downloaded all coding annotations for each of the species used in the phylogenetic analysis from GenBank as well as two additional taxa: *Vibrio cholerae* (NC_002506) and *V. parahaemolyticus* (NC_004603). We created annotations of the *Sodalis* genomes assembled here by uploading them to the RAST server (*84*). Orthologous groups were determined using Proteinortho (*85*) with a minimum connectivity of 0.8. These orthologous groups were aligned with Prank (*86*) and gene trees were inferred for all loci represented by at least four taxa using RAxML v7.3.0 and the GTRGAMMA model of nucleotide substitution. This set of gene trees was examined using ASTRAL (*87*–*89*).

Once the most likely species tree was determined, we repeated the phylogeny inference procedure used above (Supplementary Information). However, this time the Mauve alignment included the four *Sodalis* genomes assembled here and the inferred phylogeny was constrained to the topology inferred from the gene tree inference.

### Identifying coinfections

The aligned set of consensus genomes was examined to identify SNP’s fixed between the two *Sodalis* lineages assembled from *L. albipes*. We required that genotypes were available for all 36 *L. albipes* individuals and that no gaps or unknown bases were present 100 bases up- or downstream for a SNP to be used. In order to get an accurate estimate of the frequency of coinfections by both lineages, we examined individual read pairs for the presence of these SNPs. Read pairs were only counted as representing a particular lineage if at least two SNPs were present and consistent with a single lineage and there were no conflicting genotypes. We only examined read pairs mapped to a single site with no supplementary alignments and that did not require insertions or deletions to be mapped. We also excluded SNP sites that had coverage more than twice the standard deviation greater than or less than the mean coverage at these SNPs.

### Relaxed selection

To quantify the degree of relaxed selection in the *Sodalis* genomes assembled here, we used the free ratios model in PAML v4.9 (*90*, *91*) to estimate dN/dS ratios for all orthologous groups represented by at least four taxa. We required that aligned loci be at least 300 bases long, that dN/dS ratios be estimated as less than 20 and greater than 0.0001 and that the estimated dS value be at least 0.001. These quality controls were particularly important given that several of the terminal branch lengths are quite short. Statistical differences between distributions of dN/dS ratios were calculated using Wilcoxon rank-sum tests. We also had interest in determining the degree of relaxed selection in previously discovered and potentially obligate endosymbionts of honey bees. We, therefore, examined three honey bee endosymbionts with fully sequenced genomes and compared each of them to three of their close relatives using the same methods (Supplementary Information).

### *Sodalis* genome function

Genes were assigned subsystem functions using the RAST server. The counts of genes assigned to each subsystem for the free-living *S. praecaptivus* genome were used as the baseline expected number of genes. Each newly identified *Sodalis* lineage was compared to this baseline.

### PCR localization

We attempted to localize *Sodalis* in halictid bodies by extracting DNA from five body parts (head, thorax, abdomen, antennae, legs) of three male *L. albipes* and attempting to amplify this *Sodalis* locus from each of these parts (Supplementary Information). DNA extractions were done using the Qiagen Blood & Tissue DNeasy kit (Qiagen, Hilden, Germany) and results of the PCR were visualized using gel electrophoresis. We ran PCR’s of the COI barcoding locus using the primers Jerry and Pat (Magnacca & Danforth 2006) on all of the same DNA extractions to confirm DNA quality. All three negative control PCR’s that included all reagents but no DNA extract yielded no PCR product.

## Acknowledgements

We thank C. Baker, L. Henry, and J. Russell for providing insightful comments on earlier versions of this manuscript. C. Moreau, L. Pallares, A. Sedghifar, and Princeton’s EvoGroup provided valuable feedback and discussion. We also thank E. Colbert, A. Finklestein, J. Squires, L. Tomkinson, and J. Couget for their assistance with sample collection and preparation.

## Supplementary figures

**Figure S1.**

Rarefaction curves for all halictids examined in this study.

**Figure S2.**

Sites from which samples were collected. Barplots show the average bacterial community composition for social and solitary samples collected from each site. The three most common bacterial taxa are colored and all other taxa are shown in grayscale. Histograms show the proportions of shotgun sequencing reads from *L. albipes* samples from six locations (Calais, Dordogne, Rimont, Vosges, Brassus, and Ventoux) that mapped to *Sodalis*.

**Figure S3.**

Boxplots show the quantity of bacteria present in each sample as determined by qPCR of the bacterial 16S rRNA gene. Polymorphic taxa are colored by the behavior of the samples included.

**Figure S4.**

Heatmap of the presence or absence of *Sodalis* classified OTUs that appear in at least 5 samples in all *Lasioglossum* used in the comparison of social and solitary samples.

**Figure S5.**

Heatmap of the presence or absence of *Wolbachia* and *Rickettsia* classified OTUs that appear in at least five individuals in the five *Lasioglossum* species used in the between-species comparisons. Species boundaries are somewhat apparent based on the presence of these bacterial taxa.

**Figure S6.**

Proportion of genes belonging to each functional subsystem in each of the *Sodalis* lineages discovered in halictids relative to the number of genes identified in the closely-related free-living *S. praecaptivus*. Subsystem membership was determined using RAST and numbers of genes classified to each function in *S. praecaptivus* are given in parentheses.

**Supplementary tables Table S1.**

Amplicon sequencing sample information.

**Table S2.**

Full OTU table.

**Table S3.**

Statistical comparisons of core taxa in social and solitary samples.

**Table S4.**

Summary of all supervised learning analyses conducted.

**Table S5.**

Shotgun sequencing sample information.

